# N-phosphonacetyl-L-aspartate enhances type I interferon anti-viral responses through activation of non-canonical NOD2 signaling

**DOI:** 10.1101/2022.02.08.479597

**Authors:** András K. Ponti, Megan T. Zangara, Christine M. O’Connor, Erin E. Johnson, Christine McDonald

## Abstract

Type I interferon production and the expression of interferon-stimulated genes (ISGs) are key components of an innate immune response to many microbial pathogens. Dysregulation of this response can result in uncontrolled infection, inflammation, and autoimmune disease. Understanding the molecular mechanisms shaping the strength of type I interferon signaling may provide critical insights into infection control strategies and autoimmune disease therapies. Nucleotide-binding oligomerization domain 2 (NOD2) is an intracellular pattern recognition receptor that acts as both a bacterial sensor protein and a mediator of antiviral responses. Antibacterial functions of NOD2 are enhanced by treatment with the small molecule inhibitor of pyrimidine biosynthesis *N*-phosphonacetyl-L-aspartate (PALA), though how this might function in the host antiviral response remains unknown. Therefore, we tested the ability of PALA to enhance NOD2-dependent antiviral responses. Alone, PALA treatment of macrophages was not sufficient to induce interferon β (IFNβ) production or ISG expression. Instead, PALA synergized with IFNβ stimulation to enhance expression and activation of interferon-stimulated gene factor 3 (ISGF3) and induce the upregulation of a subset of ISGs in co-treated cells. Furthermore, PALA treatment of epithelial cells resulted in impaired viral replication of the herpesvirus, human cytomegalovirus. Induction of the PALA-enhanced antiviral response required activation of non-canonical NOD2 signaling mediated by mitochondrial antiviral-signaling protein (MAVS) and interferon response factor 1 (IRF1), rather than the classical receptor-interacting serine/threonine protein kinase 2 (RIP2) pathway or other IRFs previously reported to mediate NOD2 antiviral responses. These findings highlight pyrimidine metabolism enzymes as controllers of antimicrobial responses and suggest novel mechanisms for the modulation of type I interferon responses and antiviral activity.

**Significance Statement:** Understanding the molecular mechanisms shaping the strength of type I interferon signaling may provide critical insights to improve infection control strategies and autoimmune disease therapies. This work demonstrates that the pyrimidine synthesis inhibitor *N*-phosphonacetyl-L-aspartate synergizes with type I interferon to enhance antiviral responses through activation of a non-canonical NOD2 signaling pathway. These findings highlight pyrimidine metabolism enzymes as controllers of antimicrobial responses and suggest novel mechanisms for the modulation of type I interferon responses.

## Introduction

Type I interferon (IFN-I) production and the expression of interferon-stimulated genes (ISGs) are key components of an innate immune response to many microbial pathogens (1). The interferon family is comprised of 14 IFNα subtypes, IFNβ, IFNω, IFNε, IFNĸ, IFNζ, and IFNτ, and is widely recognized to induce both an antiviral response within infected cells, as well as promoting the upregulation of protective molecules in neighboring cells and systemically. IFN-I also shapes the recruitment and effector functions of immune cells in response to infection; acting on epithelial cells and tissue resident immune cells to promote the recruitment of natural killer (NK) cells, inflammatory monocytes, and dendritic cells to sites of infection where IFN-I can further enhance NK cell, dendritic cell, and T-cell functions. In addition, the immune stimulatory actions of IFN-I shape the response to bacterial infections (2) and development of cancer immunosurveillance (3). However, activation of IFN-I responses are not universally beneficial to the host and can result in uncontrolled bacterial infection, tissue damage, and autoinflammatory diseases, such as systemic lupus erythematosus and rheumatoid arthritis (4). Therefore, understanding the molecular underpinnings of the IFN-I response may have significant impact on approaches to infection control, cancer immunotherapies, and treatment of autoinflammatory diseases.

IFN-I is produced by cells in response to the sensing of viral, bacterial, or cell damage signals by pattern recognition receptors (PRRs) (1). PRRs include members of the transmembrane Toll-like receptor family (TLRs) and cytoplasmic RIG-I receptors (RLRs), nucleotide-binding oligomerization domain receptors (NLRs), and DNA sensors (e.g. cGAS-STING). Downstream activation of interferon regulator factors (IRF1, IRF3, IRF5, and IRF7) result in transcriptional induction of IFN-I and an initial wave of ISGs. IFN-I is then sensed by cells in an autocrine, paracrine, and systemic manner through binding to the cell surface receptors, IFNAR-I and IFNAR-II, stimulating the activation of Janus kinases and signal transducer and activators of transcription (STAT) complexes. The canonical STAT complex activated by IFN-I is interferon stimulated gene factor 3 (ISGF3) that is comprised of STAT1, STAT2, and IRF9. The binding of ISGF3 to specific DNA sequences called interferon stimulated response elements (ISREs) activates the transcription of a broad array of ISGs to induce an antiviral state, upregulate immunomodulatory molecules, as well as further amplify IFN-I signaling.

One PRR known to induce IFNβ production is nucleotide-binding oligomerization domain 2 (NOD2) (5). NOD2 is best known for its role as a cytoplasmic bacterial sensor protein, but also mediates antiviral responses to ssRNA and dsDNA viruses, such as respiratory syncytial virus (RSV) (6) and human cytomegalovirus (HCMV) (7, 8). Classically, NOD2 senses the peptidoglycan fragment muramyl dipeptide (MDP) component of the bacterial cell wall and triggers a signaling cascade initiated by receptor-interacting serine/threonine protein kinase 2 (RIP2), resulting in the activation NFĸB, AP-1, and IRF5 transcription factors. Activation of IRF5 by NOD2 is essential for the induction of *Ifn*β transcription in response to *Mycobacterium tuberculosis* infection (9). IFNβ production can also be stimulated as part of an antiviral response to RSV infection via the formation of a protein complex containing NOD2 and mitochondrial antiviral-signaling protein (MAVS) that results in the activation of IRF3 (6).

Our prior work demonstrated that NOD2 interacts with the pyrimidine synthesis enzyme carbamoyl-phosphate synthetase 2/ aspartate transcarbamylase/ dihydroorotase (CAD) and that CAD negatively regulates NOD2 antibacterial activity (10). Conversely, NOD2 antibacterial activity is enhanced by the CAD inhibitor *N*-phosphonacetyl-L-aspartate (PALA) (10, 11). Interestingly, other pyrimidine synthesis inhibitors have been developed to target this metabolic pathway, and many stimulate broad-spectrum antiviral activity in addition to suppressing nucleotide synthesis (12). Therefore, we investigated whether PALA modulates IFN-I production through activation of NOD2 antiviral signaling in macrophages. Similar to other pyrimidine synthesis inhibitors, we observed that PALA amplified the production of IFNβ in macrophages treated with IFN-I. However, unlike inhibitors targeting other pyrimidine synthesis pathway enzymes, the PALA-enhanced IFN-I response resulted in upregulated expression and prolonged activation of ISGF3. The PALA/IFN-I amplified response required activation of non-canonical NOD2 signaling mediated by MAVS and IRF1, rather than the classical RIP2 pathway or other IRFs previously reported to mediate NOD2 antiviral responses. These findings uncover a novel signaling pathway that modulates IFN-I responses.

## Results

### Inhibition of enzymes governing pyrimidine synthesis enhances the type I interferon response via distinct mechanisms

*De novo* pyrimidine synthesis is a multistep process requiring six different enzymatic activities housed in three proteins to generate pyrimidines (13) (SI Appendix, Fig. S1). Pharmacological inhibitors have been developed to target each enzyme of this metabolic pathway and many of these drugs also stimulate broad-spectrum antiviral immune responses; however, it is unclear whether these compounds induce an antiviral state using the same mechanism of action (12). Therefore, we compared the induction of IFNβ in response to inhibitors of CAD or dihydroorotate dehydrogenase (DHODH), as well as the ability of these inhibitors to enhance IFN-I signaling in Raw264.7 macrophages. Inhibitors that target the carbamoyl phosphate synthetase activity of CAD (acivicin), the aspartate transcarbamylase activity of CAD (PALA), or the dihydroorotase dehydrogenase activity of DHODH (brequinar and leflunomide metabolite A77 1726; teriflunomide) were tested in these experiments. Raw264.7 cells were stimulated with IFNβ alone, the inhibitor alone, or a combination of IFNβ and inhibitor for 24h and levels of secreted IFNβ quantified by ELISA. In accordance with previous reports (14, 15), all the pyrimidine synthesis inhibitors tested did not stimulate IFNβ production alone, but instead amplified IFN-I-induced IFNβ secretion (Fig. 1A-D). As these drugs target different enzymatic activities, these results suggest that amplification of IFNβ production is a general response to pyrimidine synthesis inhibition.

**Figure 1. :**
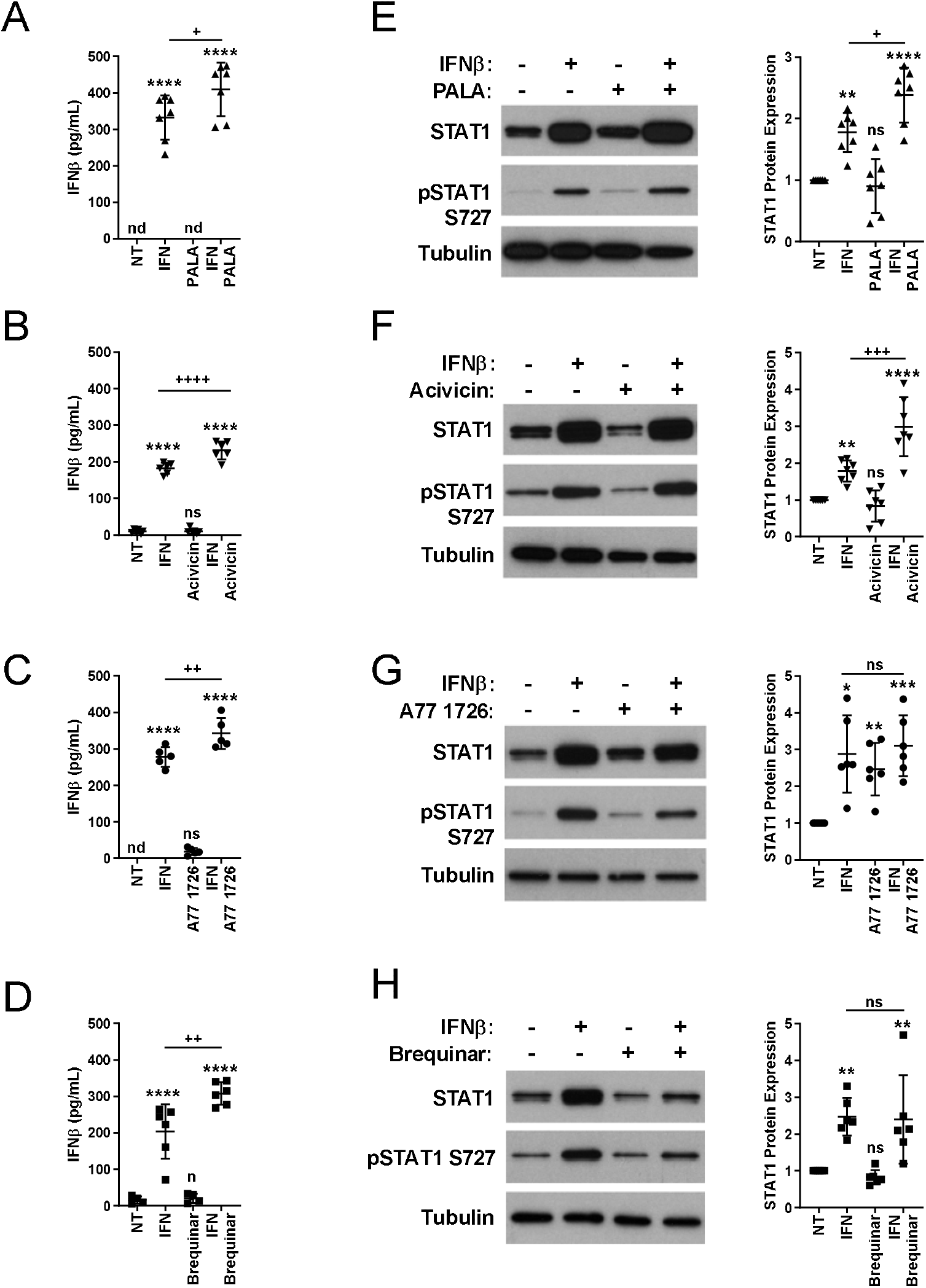
Inhibition of enzymes governing pyrimidine synthesis enhances the type I interferon response via distinct mechanisms. **A-D.** Pyrimidine synthesis inhibitors enhance IFNβ secretion. Raw264.7 cells were left unstimulated (NT), stimulated with IFNβ (200u/mL; IFN), PALA (100μM), acivicin (10μM), A77 1726 (10μM), brequinar (1μM), or a combination of IFN and pyrimidine synthesis inhibitor for 24h. IFNβ secretion was measured from cell supernatants by ELISA (n=5-7 separate experiments assayed in triplicate). Data represented as mean±SEM; significance determined by one-way ANOVA and Tukey’s multiple comparisons test; nd=not detected, ns=not significant, **** p≤0.0001 vs. NT, + p≤0.05. ++ p≤0.01, ++++ p≤0.0001 vs. IFN. **E.-H.** CAD inhibitors enhance STAT1 protein expression and prolong STAT1 activation. Raw264.7 cells were treated as in A.-D. and cell lysates analyzed by immunoblot with anti-STAT1 or anti-STAT1 phospho-S727 antibodies (left). Tubulin served as the loading control. Levels of STAT1 protein were quantified by densitometry and shown as mean±SEM (n=6-7 separate experiments); significance determined by one-way ANOVA and Tukey’s multiple comparisons test; ns=not significant, * p≤0.05, ** p≤0.01, *** p≤0.001, **** p≤0.0001 vs. NT, + p≤0.05, +++ p≤0.001 vs. IFN.

Analyses of the molecular pathways activated by pyrimidine synthesis inhibitors that mediate the enhancement of the IFN-I response revealed that inhibiting CAD and DHOH exert their effects via different mechanisms. To determine the impact of the CAD and DHODH inhibitors on STAT protein expression and activation, we treated Raw264.7 cells with IFNβ with or without inhibitors for 24h. This resulted in upregulated STAT1 expression, and additionally, IFNβ-induced STAT1 expression was further increased by co-treatment with the CAD inhibitors, PALA or acivicin (Figs. 1E, 1F), but not by DHODH inhibitors (Figs. 1G, 1H). Furthermore, PALA or acivicin treatment resulted in sustained IFNβ-stimulated phosphorylation of STAT1 at S727, indicating prolonged STAT1 transcriptional activation (Figs. 1E, 1F). Prolonged STAT1 phosphorylation of the IFN-I response was not observed in response to DHOH inhibition (Figs. 1G, 1H). These findings indicate that even though they are components of the same metabolic pathway, CAD modulates the IFN-I response using molecular mechanisms distinct from DHODH.

### Treatment with the CAD inhibitor PALA enhances ISGF3 expression and activation in response to IFN-I

IFN-I stimulates gene transcription through the activation of the ISGF3 complex comprised of STAT1, STAT2, and IRF9. As both CAD inhibitors, acivicin and PALA, enhanced the activation of the IFN-I response via STAT1, we assessed the impact of PALA treatment on the expression and activation of ISGF3 in response to IFN-I. IFNβ-induced transcription of *Stat1* and *Stat2* after 2h of treatment and it was amplified by co-treatment with PALA (Figs. 2A, 2B). This resulted in increased protein expression of all ISGF3 components at 24h, with IFNβ-stimulated expression of STAT1, STAT2, and IRF9 all increased by PALA co-treatment (Fig. 2C). The enhanced expression of STAT1 occurs at the transcriptional level, as stimulation of fibroblasts expressing STAT1 cDNA from an exogenous promoter did not result in upregulation of STAT1 protein levels (SI Appendix, Fig. S2). PALA stimulation alone was not sufficient to increase either the transcript or protein levels of ISGF3 components (Fig. 2), suggesting that PALA synergizes with IFNβ to upregulate ISGF3 transcription.

**Figure 2.**
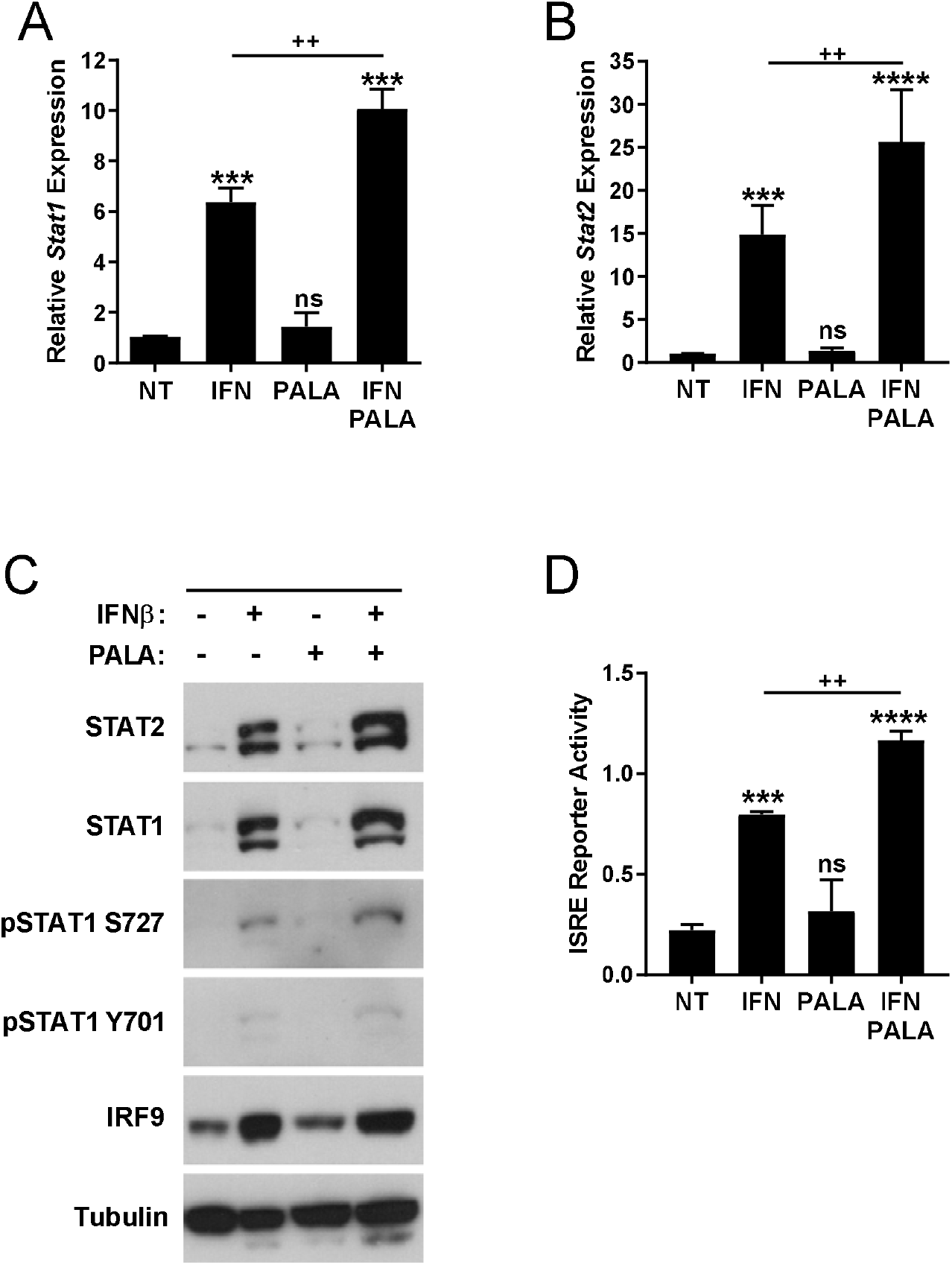
Treatment with the CAD inhibitor PALA enhances ISGF3 expression and activation in response to type I interferon. **A. & B.** Transcription of STAT1 and STAT2 in response to IFNβ is enhanced by PALA co-treatment. qRT-PCR analysis of *Stat1* (**A.**) or *Stat2* (**B.**) transcript levels from Raw264.7 cells stimulated with IFNβ (200u/mL), PALA (100uM), or co-stimulation with IFNβ and PALA for 2h (n=4). ns=not significant, *** p≤0.001, ++ p≤0.01. **C.** IFNβ-stimulated expression of ISGF3 proteins (e.g. STAT1, STAT2, IRF9) is further enhanced by PALA co-treatment. Lysates collected after 24h stimulation with IFNβ, PALA, or co-stimulation with IFNβ and PALA were probed with anti-STAT1, anti-STAT2, or anti-IRF9 to determine ISGF3 protein levels and anti-STAT1 phospho-S727 and anti-STAT1 phospho-Y701 were used to examine ISGF3 activation. Tubulin served as the loading control. Representative blots of 3 independent experiments. **D.** IFNβ-stimulated ISRE-mediated gene transcription is increased by PALA co-treatment. B16-Blue IFNα/β cells were treated with IFNβ, PALA, or IFNβ and PALA for 24h. SEAP levels in the culture supernatant were detected by colorimetric enzyme assay (n=3 experiments assayed in triplicate). Data presented as mean±SEM; significance determined by one-way ANOVA and Tukey’s multiple comparisons test; ns=not significant, *** p≤0.001, ****p≤0.001 vs. NT, ++ p≤0.01 vs. IFN.

We also found that PALA co-treatment with IFN-I correlated with increased ISGF3 expression and activation. Our data reveal that PALA co-treatment increased IFNβ-stimulated STAT1 phosphorylation at both serine 727 (S727) and tyrosine 701 (Y701) sites, which are indicative of STAT1 activation (Fig. 2C). Activation of the ISGF3 complex was measured using a reporter gene assay, where expression of secreted embryonic alkaline phosphatase (SEAP) is driven by interferon stimulated response elements (ISRE). While the ISRE reporter gene was not activated by PALA treatment alone, co-treatment of the reporter cells with IFNβ and PALA enhanced expression of the ISRE-driven reporter gene over the levels induced by IFNβ alone (Fig. 2D). These results indicate that the combined treatment of PALA with IFN-I upregulates both ISGF3 expression and activation of this transcription factor complex.

### PALA amplifies a subset of IFNβ-stimulated transcripts and induces an anti-viral state

We next assessed the impact of PALA and IFN-I on transcription of IFN-stimulated genes. Co-treatment of cells with PALA and IFNβ resulted in a significant enhancement of ISG expression that included *Mx1, Isg15, Ifit1*, and *Ifit2* (Figs. 3A, 3B). In contrast to previous reports in which ISGs were induced by DHODH inhibitors (16), *Oas1* transcription was not enhanced by PALA co-treatment of IFNβ-stimulated cells, but instead were suppressed (Fig. 3B). These findings demonstrate that PALA induces a unique set of target genes that only partially overlaps with the reported mechanism of antiviral induction by DHODH inhibitors.

**Figure 3:**
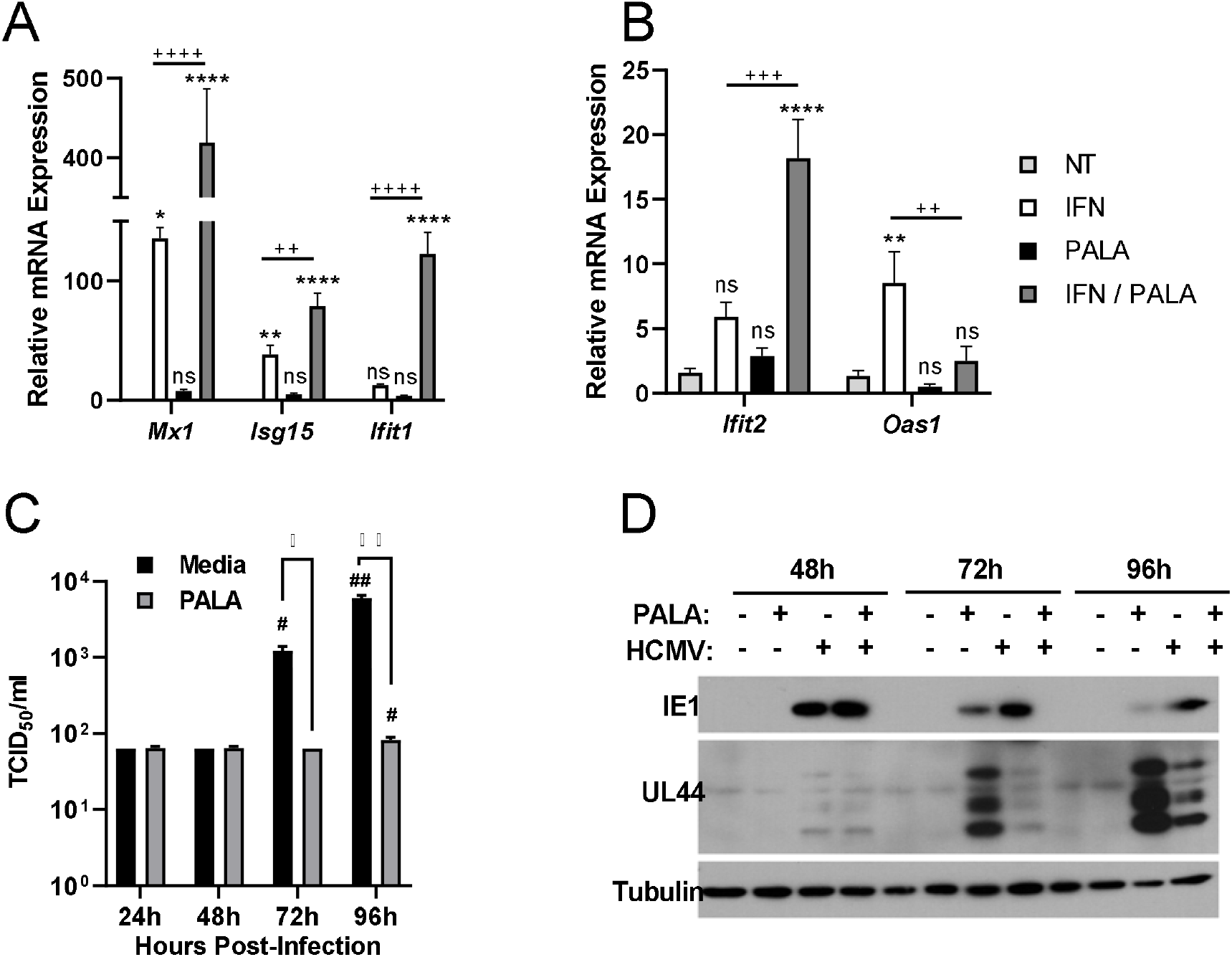
PALA enhances the induction of an antiviral state by type I interferon. **A. & B.** Analysis of the impact of PALA co-treatment on ISG transcripts induced by IFNβ by qRT-PCR. *Mx1, Isg15, and Ifit1* (**A.**) or *Ifit2* and *Oas1* (**B.**) transcript levels in Raw264.7 cells treated with IFNβ (200u/mL), PALA (100uM), or co-stimulation with IFNβ and PALA for 24h (n=4 experiments). Data presented as mean±SEM; significance determined by one-way ANOVA and Tukey’s multiple comparisons test; ns=not significant, *** p≤0.001 vs. NT, ++ p≤0.01 vs. IFN. **C.** PALA suppresses HCMV replication. ARPE-19 cells were infected with HCMV (MOI=3 TCID_50_/cell) and then cultured in media ±PALA (100μM). *De novo* cell-associated virus production was titered by TCID_50_ assay on naïve NuFF-1 cells (n=3 experiments performed in triplicate; representative experiment shown). Data presented as mean±SEM; significance determined by one-way ANOVA and Sidak’s multiple comparisons test; ns=not significant, *p≤0.05, **p≤0.01 media vs. PALA; #p≤0.05, ##p≤0.01 vs. 24h sample. **D.** ARPE-19 cells were infected (MOI=3 TCID_50_/cell) and lysates collected at the indicated times post-infection. Expression of viral proteins were assessed by immunoblot probed with anti-IE1, anti-pUL44, and anti-tubulin (n=3 experiments; representative experiment shown).

We next assessed the ability of PALA to induce an antiviral response using the dsDNA herpesvirus, human cytomegalovirus (HCMV), as a model. Upregulation of the IFN-I response following HCMV infection limits viral replication in an IFNα receptor-dependent manner as part of the innate antiviral response (17). Human retinal pigmented epithelial cells (ARPE-19) were infected with HCMV (MOI=3 TCID_50_/cell) and then treated with or without PALA over a 96h time course and HCMV replication was monitored by tissue culture infectious dose (TCID_50_) assay. PALA significantly suppressed HCMV infectious titers at 72h and 96h post-infection (Fig. 3C). To determine the stage of infection at which PALA impacts HCMV replication, we assessed the accumulation of viral proteins representative of immediate early responses (IE) and early (E) kinetic classes, IE1 and pUL44 respectively, over a 96h time course. The expression of IE genes and their translated proteins induce expression of early viral genes, whose proteins are required for viral DNA replication (18, 19). PALA treatment resulted in higher accumulation of IE1 at all time points, but markedly reduced levels of pUL44 at 72h and 96h post-infection (Fig. 3D). This indicates that PALA limits translation of early proteins, resulting in repression of viral replication independent of cellular pyrimidine depletion. Collectively, these data demonstrate that PALA, under conditions that stimulate an IFN-I response, effectively induces an antiviral state.

### PALA promotes increased antiviral responses through de-repression of NOD2 signaling

CAD binds to and negatively regulates the antibacterial activity of the intracellular pattern recognition receptor, NOD2 (10), thus demonstrating a direct link between pyrimidine synthesis and innate immune modulation. We tested whether the enhancement of IFN-I responses by PALA co-treatment was dependent on NOD2 expression. Bone marrow-derived macrophages (BMDMs) were isolated from C57BL/6 (WT) or *Nod2^−/−^* (NOD2KO) mice and stimulated with IFNβ, PALA, or a combination of IFNβ and PALA for 24h and STAT1 expression levels were determined by immunoblot. Although IFNβ stimulated STAT1 protein expression in both WT and NOD2KO macrophages, NOD2KO macrophages were unresponsive to PALA treatment (Fig. 4A). These results indicate that NOD2 expression is required to mediate PALA enhancement of a type I IFN response.

**Figure 4:**
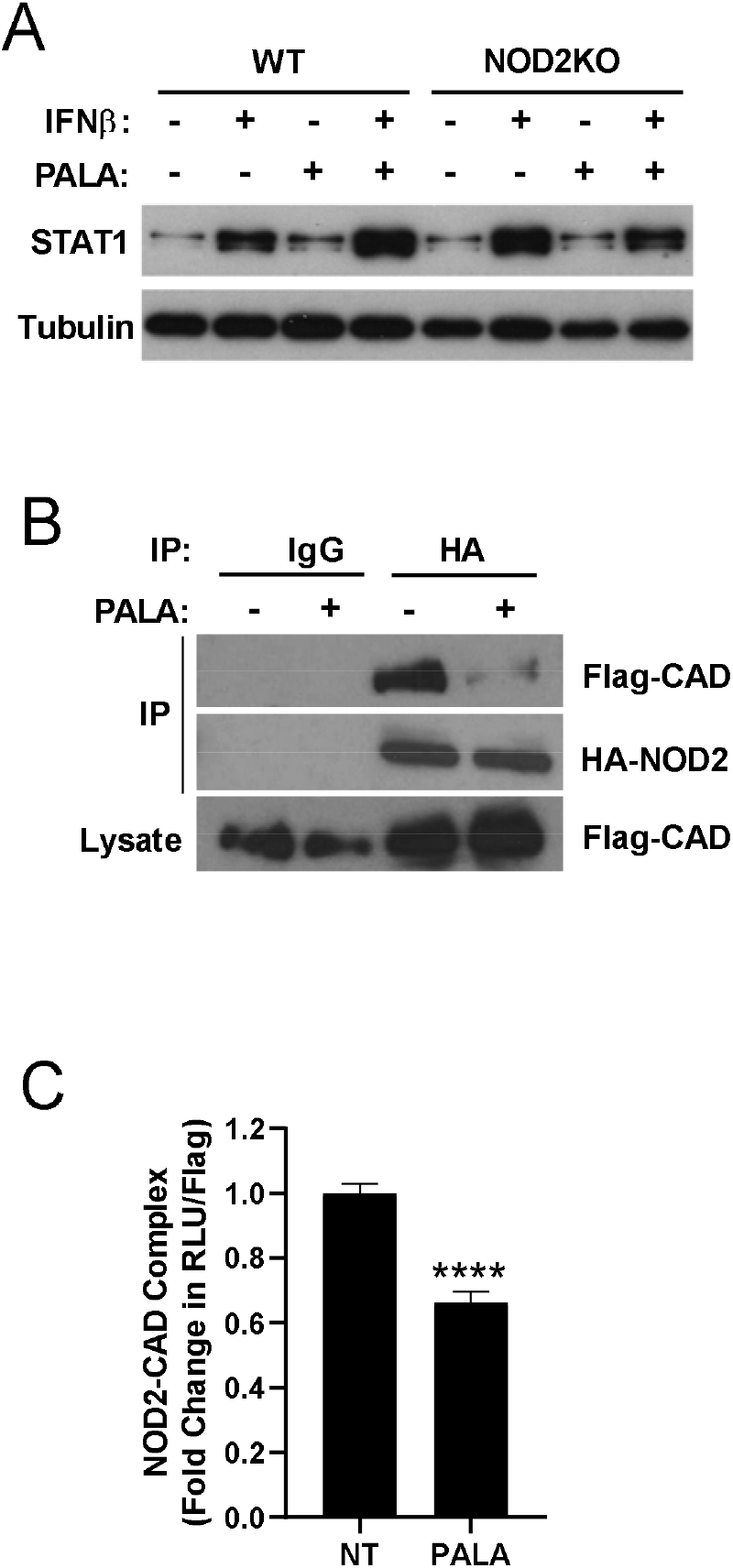
PALA promotes increased antiviral responses through de-repression of NOD2 signaling. **A.** PALA-enhanced STAT1 protein levels requires NOD2 expression. Immunoblot analysis of STAT1 protein levels in BMDM from C57BL/6 (WT) or *Nod2^−/−^* (NOD2KO) mice stimulated with IFNβ (200u/mL), PALA (100uM), or co-stimulation with IFNβ and PALA for 24h (n=3). Tubulin served as a loading control. **B.** PALA induces dissociation of a NOD2-CAD protein complex. HEK293T cells were transfected with plasmids expressing Flag-tagged CAD (Flag-CAD) and HA-tagged NOD2 (HA-NOD2) and treated ± PALA (50μM; 2h). Cell lysates were immunoprecipitated (IP) with mouse IgG or anti-HA antibodies and proteins visualized by immunoblot with anti-Flag or anti-HA antibodies. Image representative of 4 separate experiments. **C.** PALA treatment disrupts a CAD-NOD2 protein complex. HEK293T cells expressing Renilla-tagged NOD2 and Flag-tagged CAD constructs were treated ± PALA (100μM; 2h). Lysates were incubated with plate-bound anti-FLAG antibody and luciferase measured (RLU) to quantitate the amount of co-precipitating NOD2. The amount of plate-bound CAD was determined by anti-Flag ELISA and luciferase values normalized to CAD levels. Data presented as mean±SEM (n=6 experiments); significance determined by Student’s t-test, **** p≤0.0001

NOD2 forms a protein-protein complex with CAD (10) and knockdown of CAD expression or PALA treatment enhances NOD2 activity (10, 11). Therefore, we next investigated whether PALA relieves CAD inhibition of NOD2 activity through disruption of the CAD-NOD2 protein complex. PALA treatment of HEK293T cells expressing epitope-tagged CAD and NOD2 proteins decreased the amount of CAD co-immunoprecipitating with NOD2 (Fig. 4B). We also quantitatively assessed the PALA-induced disruption of the CAD-NOD2 complex using the Lumier with Bacon assay (20) that measures the amount of a Renilla-tagged NOD2 associating with a quantified amount of plate-bound Flag-CAD through sequential luminescent and ELISA readings of the same well. As in Fig. 4B, we found a reduction in co-immunoprecipitation of NOD2 by CAD (Fig. 4C), which correlated to a significant reduction in luciferase levels in PALA treated samples (~35% decrease) by the Lumier with Bacon assay (Fig. 4C). These results suggest PALA disrupts an inhibitory CAD-NOD2 protein complex that results in NOD2-mediated enhancement of IFN-I responses.

### Induction of the PALA enhanced antiviral response requires activation of non-canonical NOD2 signaling mediated by mitochondrial antiviral-signaling protein (MAVS)

NOD2 activates the production of IFN-I through a signaling cascade dependent on RIP2 (21), as well as a pathway mediated by MAVS (6), depending on the NOD2 activating signal. Therefore, we assessed the involvement of known mechanisms of NOD2 innate immune signaling in the PALA enhancement of IFN-I responses using knockout and RNAi knockdown cells. First, we tested the requirement of RIP2 in the PALA enhancement of an IFN-I response by assaying the upregulation of STAT1 protein levels by immunoblot of lysates of BMDM from WT or *Ripk2^−/−^* (RIP2KO) mice. Although IFNβ increased STAT1 protein levels in both WT and RIP2KO BMDM, RIP2 expression was not required for upregulation of IFNβ-stimulated STAT1 by PALA co-treatment (Fig. 5A). Next, we evaluated the requirement of non-canonical NOD2 signaling via MAVS in Raw264.7 cells treated with MAVS-targeting RNAi. MAVS knockdown resulted in the loss of increased STAT1 protein expression in response to co-treatment with PALA and IFNβ (Fig. 5B). Complementary to this finding, enhanced ISRE-mediated transcription induced by PALA and IFNβ co-treatment was blocked in B16-Blue IFNα/β reporter cells treated with MAVS-targeting RNAi (Fig. 5C), suggesting that CAD inhibition promotes NOD2 signaling through MAVS to enhance the IFN-I response.

**Figure 5:**
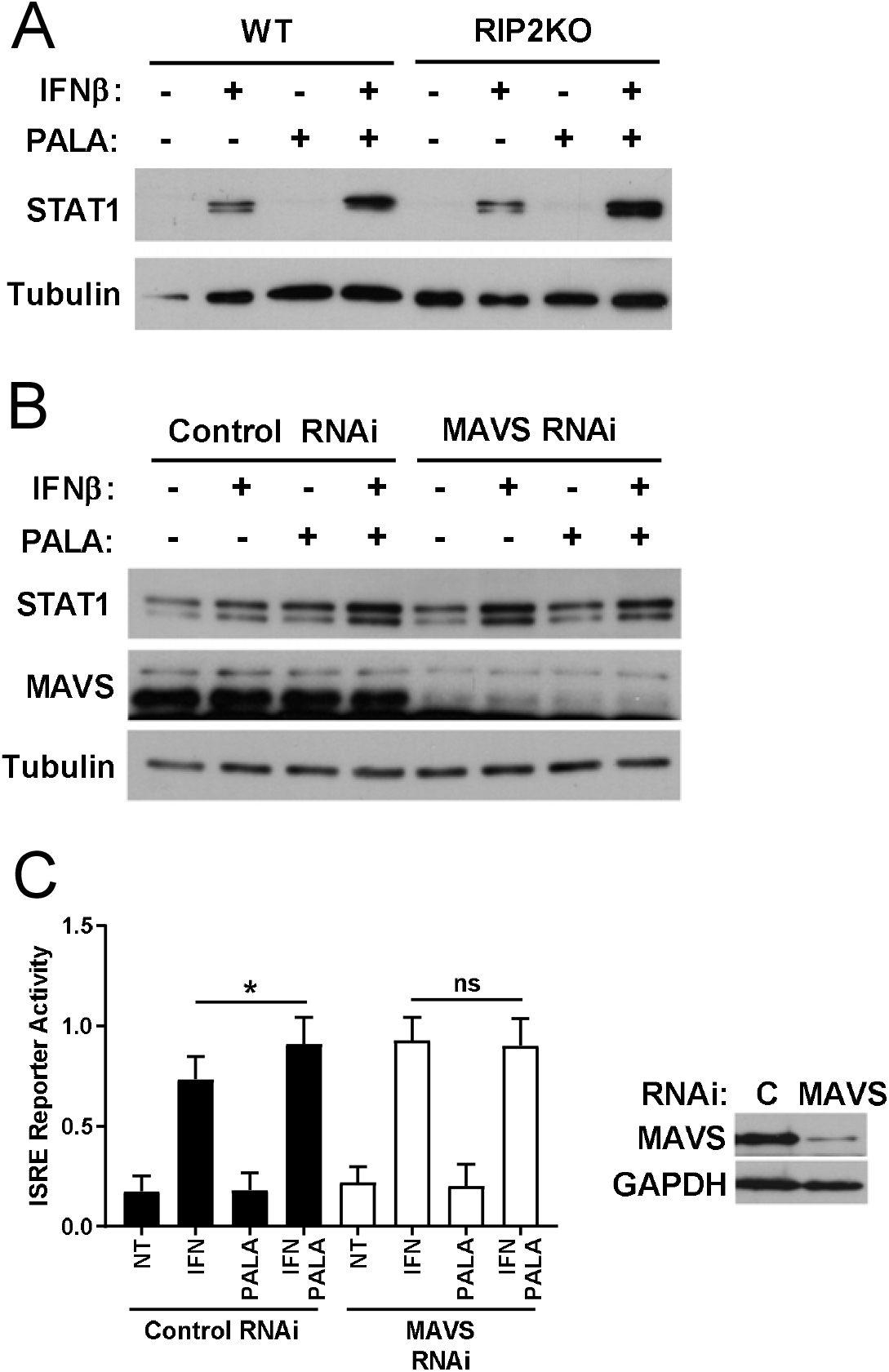
Induction of the PALA enhanced antiviral response requires activation of non-canonical NOD2 signaling mediated by MAVS. **A.** RIP2 expression is not required for PALA enhancement of STAT1 protein levels. STAT1 protein levels in BMDM from C57BL/6 (WT) or *Rip2^−/−^* (RIP2KO) mice stimulated with IFNβ (200u/mL), PALA (100uM), or co-stimulation with IFNβ and PALA for 24h. Representative immunoblot from 4 independent experiments shown. **B.** MAVS expression is required for upregulation of IFNβ-stimulated STAT1 expression by PALA. Raw264.7 cells were transfected with non-targeting (Control) or MAVS-targeting RNAi for 72h. Cells were treated as in A. and analyzed for STAT1, MAVS, and tubulin protein levels (n=4). **C.** B16-Blue IFNα/β reporter cells were transfected with non-targeting (Control) or MAVS-targeting RNAi for 72h. Cells were treated as in A. and ISRE reporter assayed through quantification of SEAP levels in the media by colormetric assay (n=4 performed in triplicate). Data presented as mean±SEM; significance determined by one-way ANOVA and Tukey’s multiple comparisons test; ns=not significant, * p≤0.05 vs. IFN. Inset shows MAVS and tubulin protein levels to confirm RNAi-mediated knockdown of expression.

### PALA induces activation of a non-canonical NOD2 signaling pathway mediated by IRF1

It is accepted that IRFs signal downstream of MAVS as part of the IFN-I response to viruses (22). For example, the NOD2-MAVS-IRF3 axis mediates an IFN-I response to ssRNA and respiratory syncytial virus (RSV) infection (6). Surprisingly, PALA did not impact the activation and nuclear translocation of IRF3 or IRF5, either alone or in combination with IFNβ as visualized in immunoblots of fractionated RAW264.7 cells (Fig. 6A). However, significantly more IRF1 localized to the nuclei of RAW264.7 cells co-stimulated with IFNβ and PALA when assessed by immunoblots of fractionated cell lysates or visualized by immunofluorescent confocal microscopy (Figs. 6A, 6B). RNAi knockdown of IRF1 abrogated PALA enhancement of IFNβ-stimulated activation of an ISRE reporter gene in B16-Blue IFNα/β reporter cells (Fig. 6C). Finally, PALA enhanced IRF1 activation required NOD2 expression, as IFNβ-stimulated nuclear translocation of IRF1 visualized by immunofluorescent confocal microscopy was not increased by PALA treatment of NOD2KO BMDM (Fig. 6D). Taken together, these data demonstrate a novel role for the pyrimidine synthesis enzyme CAD in the modulation of a non-canonical NOD2 signaling pathway mediated by MAVS and IRF1 to enhance IFN-I antiviral responses.

**Figure 6:**
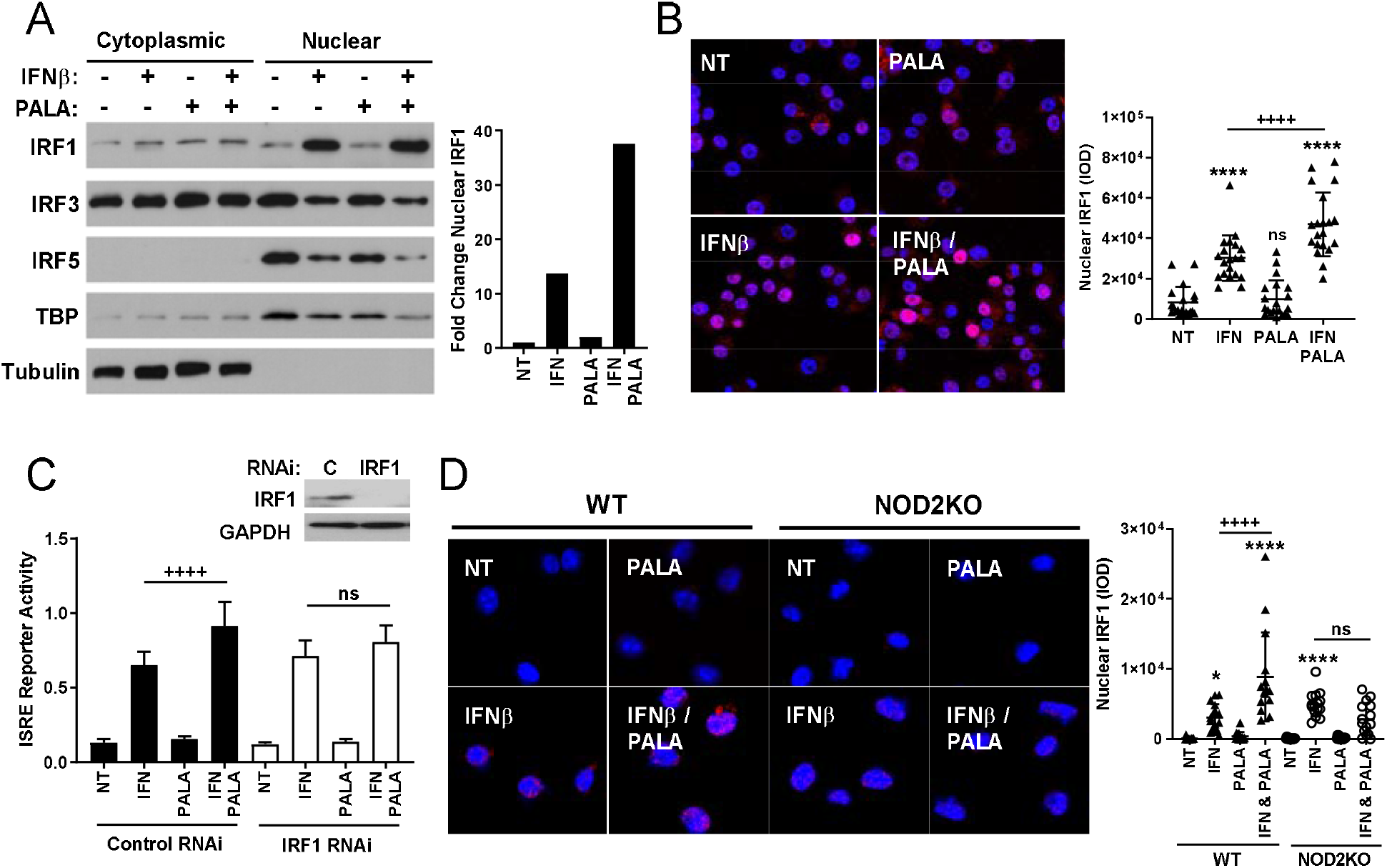
PALA induces activation of a non-canonical NOD2 signaling pathway mediated by IRF1. **A.** PALA enhances IFNβ-stimulated IRF1 nuclear localization and not activation of IRF3 or IRF5. Raw264.7 cells were cells stimulated with IFNβ (200 u/mL), PALA (100uM), or co-stimulation with IFNβ and PALA for 1h. Cell lysates were fractionated into cytoplasmic and nuclear fractions and immunoblots performed to visualize the localization of IRF1, IRF3, and IRF5 (n=3; representative experiment shown). Blots were also probed for the nuclear protein TBP and the cytosolic protein tubulin as controls. IRF1 nuclear protein levels were quantified by densitometry and fold change relative to NT (left panel). **B.** Enhancement of IFNβ activation of IRF1 by PALA can be visualized by immunofluorescent confocal microscopy. Raw264.7 cells were treated as in A., co-stained with anti-IRF1 antibody (red) and DAPI (blue), and visualized by fluorescent confocal microscopy. Intensity of IRF1 nuclear signal was quantitated using Image-Pro software. Measurements represent 5 replicate fields from 4 individual experiments. Data presented as mean±SEM; significance determined by one-way ANOVA and Tukey’s multiple comparisons test; ns=not significant, **** p≤0.0001 vs. NT, ++++ p≤0.0001 vs. IFN. **C.** IRF1 expression is required for increased IFNβ-stimulated ISRE transcription in response to PALA treatment. B16-Blue IFNα/β reporter cells were transfected with non-targeting (Control) or IRF1-targeting RNAi for 72h. Cells were treated as in A. for 24h and ISRE reporter assayed through quantification of SEAP levels in the media by colormetric assay (n=4 performed in triplicate). Data presented as mean±SEM; significance determined by one-way ANOVA and Tukey’s multiple comparisons test; ns=not significant, ++++p≤0.0001 vs. IFN. Inset shows immunoblot of IRF1 and tubulin protein levels to show RNAi-mediated knockdown of expression. **D.** Activation of IRF1 by PALA is NOD2-dependent. BMDM from C57BL/6 (WT) or *Nod2^−/−^* (NOD2KO) mice were stimulated and analyzed for IRF1 localization by immunofluorescent confocal microscopy as in B.

## Discussion

Administration of recombinant IFN-I is used clinically to control viral infections, such as hepatitis C, hepatitis B, and human immunodeficiency virus, or as antineoplastic therapies for hairy cell leukemia, melanoma, and breast cancer (23). Although IFN-I can be used as a single agent, is it is often administered with another drug, such as the nucleotide synthesis inhibitors ribavirin or gemcitabine, to increase therapeutic efficacy. Conversely, an exacerbated IFN-I signature is associated with several autoimmune and autoinflammatory diseases, including systemic lupus erythematosus (SLE), rheumatoid arthritis, and Sjören’s syndrome, as well as in chronic *Mycobacterium tuberculosis* (TB) infection, suggesting a role for this pathway in disease pathogenesis (24). Therefore, an expanded knowledge of the mechanisms underlying the induction and amplification of IFN-I driven responses may provide insights into pathogenic mechanisms of disease and new targets for therapies. Our work herein identifies a novel molecular signaling pathway regulated by the pyrimidine synthesis enzyme CAD mediated by NOD2, MAVS, and IRF1 that modulates the strength of IFN-I responses.

The signaling cascades required for IFNβ induction by NOD2 are stimulus-dependent. In response to bacterial stimuli, such as *M. tuberculosis*, NOD2 activates the protein kinase cascade of RIP2 and TANK-binding kinase 1 (TBK1) in macrophages to activate IRF5-dependent transcription of *Ifn*β (9). However, IFNβ induction in response to the ssRNA virus RSV signals through the formation of a mitochondrial associated protein complex of NOD2 and MAVS resulting in the activation of IRF3 and IRF7 (6). Finally, in response to the dsRNA mimetic, poly I:C, NOD2 forms a complex with 2’-5’-oligoadenylate synthetase type 2 (OAS2) to activate RNase-L (8). RNase-L creates RNA fragments that stimulate IFNβ production through a MDA5/IRF3 signaling pathway (8). To date, there are no reports directly linking NOD2 signaling with activation of IRF1. One report identified NOD2 and IRF1 as co-regulated transcripts in a functional screen of inflammatory bowel disease-associated loci, suggesting involvement of these proteins in similar cellular responses (25). Our results identify a novel signal transduction pathway that requires NOD2, MAVS, and IRF1 to amplify and prolong IFNβ signaling through ISGF3. This non-canonical NOD2 signaling is held in check by the pyrimidine synthesis enzyme CAD, demonstrating another mechanism of metabolic crosstalk with innate immune sensors.

Our data indicate that PALA treatment interferes with HCMV infection by suppressing expression of viral early protein synthesis in epithelial cells, suggesting that this CAD inhibitor has immune modulatory antiviral actions in addition to suppression of cellular pyrimidine synthesis. HCMV infection results in increased pyrimidine synthesis flux to provide nucleotides for genome replication and UDP-sugars for glycosylation of viral envelope proteins (26). Inhibition of pyrimidine synthesis by PALA or DHODH inhibitors successfully blocked HCMV replication and production of glycosylated envelope proteins required for infectious virions (26). However, in contrast to our findings that PALA caused a reduction in pUL44 expression, the prior study of HCMV infected embryonic lung fibroblasts showed that accumulation of viral proteins were unaffected by PALA treatment, although viral DNA abundance was substantially reduced and not rescued by addition of exogenous uridine (26). We postulate the difference between our findings and the previous work may be the higher level of basal NOD2 expression in epithelial cells versus fibroblasts. In support of this, others have shown NOD2 expression is undetectable in human foreskin fibroblasts, though HCMV infection robustly upregulates NOD2 expression in these cells (27). This also raises the possibility of tissue-specific differences in the antiviral response, as fibroblasts of different tissue origin may differentially regulate NOD2 (26, 27). Similar to our results, exogenous NOD2 expression prior to HCMV infection resulted in reduced pUL44 expression and decreased viral replication (27). These findings suggest that through NOD2 stimulation, PALA inhibits HCMV replication independent of mechanisms involving pyrimidine depletion.

Multiple nucleotide synthesis inhibitors link nucleotide biosynthesis with regulation of innate immune responses that potentiate an anti-viral response through upregulation of ISGs; however, the molecular mechanisms underlying these responses are not well-defined (12). Similar to the actions of PALA, the DHODH inhibitors, DD78 and brequinar, synergize with ssRNA stimulation to enhance production of ISGs in an IRF1-dependent manner (16). However, our data indicate the signaling pathways that mediate activation of IRF1 in response to pyrimidine synthesis inhibition may depend on which enzyme in the pathway is targeted (e.g. CAD or DHODH). While CAD and DHODH inhibitors each enhanced secretion of IFNβ, only the CAD inhibitors caused the upregulation and sustained activation of ISGF3 components. The differences in signaling intermediates are also reflected in the ISG expression profile in response to different pyrimidine synthesis inhibitors. PALA, brequinar, and DD264 all required a second signal (e.g. IFNβ or ssRNA, respectively) to shape the ISG expression profile. Both CAD and DHODH inhibitors enhanced the expression of *Isg15*, *Ifit1*, and *Ifit2*; however, opposite effects were observed for *Mx1* and *Oas1,* with PALA increasing *Mx1* expression and repressing *Oas1* transcript levels (16). Further investigation of the upstream triggers of these different response pathways is warranted to increase our understanding of how these pyrimidine synthesis inhibitors tailor expression of ISGs and antiviral responses.

Although the molecular trigger activating this non-canonical NOD2 signaling pathway has yet to be identified, we postulate that it may be a cellular stress signal, such as mitochondrial DNA (mtDNA) release into the cytosol, reactive oxygen species generation, or endoplasmic reticulum stress. NOD2 has been implicated as a cell stress sensor activated in response to endoplasmic reticulum stress or as a mediator of metabolic signaling responses triggered by fatty acids and hyperglycemia (28). The release of NOD2 from an inhibitory CAD-NOD2 complex may permit the relocation of NOD2 to sites of cell stress and triggering of an IFN-I response. For example, a major site of reactive oxygen species generation is the peroxisome and studies indicate that pattern recognition receptor engagement of MAVS localized to peroxisomes stimulates an IRF1 signaling response (29, 30). Alternately, CAD knockdown in HeLa cells resulted in transcriptional induction of ISGs that was at least partially dependent on the cytosolic release of mtDNA (31). Mitochondrial stress and mtDNA release has also been associated with a cellular IFN-I signature in molecular analyses of SLE patients and individuals with chronic TB infections (24). It will be of interest to determine in future studies if NOD2 is acting as a cell stress sensor that mediates responses to mitochondrial stress by sensing cytosolic mtDNA or metabolic stress detected at peroxisomes, especially as it may integrate cell stress with antiviral and antibacterial responses.

The utility of IFN-I treatment for emerging pandemic viral diseases, such as Ebola, severe acute repiratory syndrome (SARS), Middle East respiratory syndrome (MERS), and coronavirus disease 2019 (COVID-19), has shown promise in clinical trials (32). However, the timing of IFN-I administration is critical to the potential efficacy and avoidance of harm in these infections. In the case of SARS-CoV-2 infection, the virus that causes COVID-19, early disease is characterized by a blunted interferon response and IFN-I administration at this time may improve time to recovery. This is in contrast to later stage or in severe disease when IFN-I may enhance the cytokine storm through excessive recruitment of activated monocytes. There is interest in finding drugs that can modulate the IFN response as therapeutics for SARS-CoV-2 infection, as this approach would be potentially unaffected by emerging viral variants. In support of this is a recent study demonstrating specific DHODH inhibitors (S312 and S416) were potent antiviral agents in SARS-CoV-2 infected cells and effective in increasing the survival of mice infected with a range of RNA viruses (33). Intriguingly, S312 was effective both as a prophylactic treatment and as a therapeutic agent as late as 6 days post-infection in a mouse model of influenza A infection that, similar to COVID-19, results in a virally triggered, pro-inflammatory cytokine storm. These findings suggest additional immunomodulatory actions of pyrimidine synthesis inhibition that remain to be defined. Phase I and phase II clinical trials of DHODH inhibitors (brequinar; NCT04425252 and IMU-383; NCT04516915) have also commenced and it will be exciting to determine whether pyrimidine synthesis inhibition is an effective COVID-19 therapy.

In summary, this work demonstrates that the pyrimidine synthesis inhibitor *N*-phosphonacetyl-L-aspartate (PALA) synergizes with type I interferon to enhance antiviral responses through activation of a non-canonical NOD2 signaling pathway. This is the first description of this non-canonical pathway, comprised of NOD2, MAVS, and IRF1, that modulates IFN-I responses and expression of ISGs. Understanding the molecular mechanisms shaping the strength of IFN-I signaling may provide critical insights to improve infection control strategies and autoimmune disease therapies. These findings highlight pyrimidine metabolism enzymes as controllers of antimicrobial responses and suggest novel mechanisms for the modulation of type I interferon responses.

## Materials and Methods

### Cell Culture

RAW-264.7 (ATCC), HEK-293T (ATCC), 2fTGH (Dr. George Stark, Cleveland Clinic) and U3A-R (Dr. George Stark, Cleveland Clinic) cell lines were cultured in Dulbecco’s Modified Eagle’s medium with high glucose (DMEM) (GibcoBRL Technologies) with 10% heat-inactivated fetal bovine serum (FBS; GibcoBRL Technologies) and 2 mM L-glutamine. B16-Blue IFNα/β reporter cells (Invivogen) were cultured in RPMI 1640 medium (GibcoBRL Technologies) with 10% heat-inactivated FBS and 100 μg/mL Zeocin (InvivoGen). NuFF-1 cells (GlobalStem) were maintained in DMEM with 10% FBS, 2 mM L-glutamine, 0.1 mM non-essential amino acids, 10 mM HEPES, and 100 U/ml penicillin/streptomycin. ARPE-19 (ATCC) cells were cultured in DMEM:Ham’s F12 (1:1), with 10% FBS, 2.5 mM _L_-glutamine, 0.5 mM sodium pyruvate, 15 mM HEPES, 1.2 g/L sodium bicarbonate, and 100 U/ml penicillin/streptomycin. Cell lines were all maintained at 37°C with humidified air containing 5% CO_2_. Mouse bone marrow-derived macrophages (BMDMs) were isolated as described previously (34). Cells were treated with: recombinant human IFNβ (PeproTech), recombinant mouse IFNβ (BioLegend), PALA (Open Chemical Repository of the Developmental Therapeutics Program NCI/NIH), acivicin (Santa Cruz Biotechnology), A77 1726 (Santa Cruz Biotechnology), and brequinar (Tocris).

### Human Cytomegalovirus Infection

Stocks of the clinical isolate of human cytomegalovirus (HCMV), TB40/*Egfp*, were generated and titered by 50% tissue culture infectious dose (TCID_50_) assay on NuFF-1 cells as described previously (35, 36). To assess viral replication, ARPE-19 cells were infected at a MOI of 3 TCID_50_/cell. Viral adsorption was centrifugally enhanced for 30 min at 1000 x *g* at room temperature and then incubated for an additional 1 h at 37°C, during which the plates were rocked every 15 min. The viral inoculum was then removed, cells were washed 3 times with PBS, and cultures were replenished with fresh media in the absence or presence of 100 μM PALA. Infected cells were harvested every 24 h over a 96 h time course. At the time of harvest, cells were washed 3 times with PBS, scraped into 1 ml of fresh ARPE-19 media without PALA, and stored at −80°C. All samples were subject to 3 freeze/thaw cycles to lyse infected cells, after which the cellular debris was removed by centrifugation. The infectious media was titered by TCID_50_ assay on naïve NuFF-1 cells. Infected cells were alternatively lysed in 10 mM HEPES pH 7.4, 142 mM KCl, 5 mM MgCl_2_, 1 mM EGTA, 0.2% NP40, and 1X protease inhibitor cocktail (Sigma) and boiled with 5X Laemmli buffer for SDS-PAGE and immunoblotting.

### Immunoblotting

Whole cell lysates were prepared in 5x Laemmli buffer, separated by SDS-PAGE, and transferred to a PVDF membrane. After blocking with 5% milk/ TBST for 1 h at room temperature, membranes were probed with primary antibodies at 4°C overnight. The following Cell Signaling Technology antibodies were used: GAPDH (2118L), STAT1 (14994S), phospho-STAT1 (Ser727) (8826S), phospho-STAT1 (Tyr701) (7649S), IRF9 (28845S), IRF1 (8478), IRF5 (13496S), and TBP (8515). Additional antibodies included STAT2 (07-140, EMD Millipore), MAVS (ab31334; abcam), FLAG-M2 (F1804; Sigma), α-tubulin (T9026, Sigma), HA.11 (MMS-101P; Covance), IRF3 (655702, Biolegend), IE1 (clone 1B12) (37), and UL44 (pUL44; Virusys). Membranes were washed 3 times TBST and then incubated with donkey anti-rabbit or donkey anti-mouse HRP-conjugated secondary antibodies (Jackson ImmunoResearch Laboratories) for 1 h at room temperature. Western blots were developed using ECL reagent (EMD Millipore). Detection of the IE1 and pUL44 viral proteins used 1% BSA/TBST as a blocking agent. Western blot quantification was performed using ImageJ software (NIH).

### RNA interference (RNAi) knockdown and B16-Blue ISRE reporter assay

For RNAi knockdown of IRF1 and MAVS, B16-Blue cells were seeded at 10,000 cells/well. The following day, media was replaced with serum-free RPMI containing one of the following siRNAs: Accell Green Non-targeting siRNA (Dharmacon), Accell Mouse *Irf1* (16362) siRNA – SMARTpool (Dharmacon), and Accell Mouse Mavs (228607) siRNA – SMARTpool (Dharmacon). Cells were incubated 72 h with siRNA, then treated for 24 h with 200 U/mL IFNβ and 100 μM PALA. Media was collected and combined with QUANTI-Blue substrate (Invivogen) to assess secreted alkaline phosphatase (SEAP) levels via OD_655_.

### RNA Isolation and Quantitative RT-PCR Analysis

Total RNA was isolated from samples using the RNeasy mini kit (Qiagen) and cDNA prepared using the iScript cDNA synthesis kit (BioRad). qPCR was performed using SYBR Green supermix (BioRad) on the CFX96 Touch Real-Time PCR Detection System (BioRad). Relative *C_T_* values were calculated using the 2^-ΔΔ*Ct*^ method (38).

### ELISA Detection of IFNβ

RAW-264.7 cells were treated for 24 h with 200 U/mL IFNβ and 100 μM PALA. Secreted IFNβ levels were quantified in cell supernatants using the Mouse IFNβ, Legend Max ELISA Kit with Pre-coated Plates (BioLegend) as specified by the manufacturer.

### Immunoprecipitation

HEK-293T cells were transfected with pcDNA3, pcDNA3-HA-NOD2, and His-FLAG-CAD expression plasmids using Lipofectamine as previously described (10). After 24 h, cells were treated for 2 h with 100 μM PALA and then lysed in NP40 Lysis Buffer (10 mM HEPES ph 7.4, 142 mM KCL, 5 mM MgCl_2_, 1 mM EGTA, 0.2% NP-40, and 1X Sigma protease inhibitor cocktail). Lysates were precleared with protein G-sepharose 4B beads (Sigma) and then incubated with anti-HA antibody (HA.11; Covance) overnight at 4°C. Immunocomplexes were collected on protein G-sepharose 4B beads, washed in NP40 Lysis Buffer, and then resuspended in Laemmli Buffer for immunoblotting.

### LUMIER with BACON (39)

HEK-293T cells were seeded at 500,000 cells/well in 6 well plates. Cells were transfected the next day with either Renilla-HA-NOD2 (750 ng/well) and pcDNA3 (250 ng/well), or Renilla-HA-NOD2 (750 ng/well) and His-FLAG-CAD (250 ng/well) using Effectene Transfection Reagent (Qiagen). The following day, cells were treated with 100 μM PALA for 4 h. Cells were lysed in 1x Renilla Luciferase Assay Lysis Buffer (Promega) at room temperature with rocking. Cell lysate was added to anti-FLAG antibody pre-coated white plates in triplicate (immunoprecipitated luciferase) or to clear uncoated plates in triplicate (total luciferase) and incubated for 3 h at 4°C with rocking. Renilla luciferase signal was detected using Renilla Luciferase Assay Reagent (Promega) and a Spectramax iD3 96 well plate reader (Molecular Devices). After luciferase levels were measured, the total amount of bound His-FLAG-CAD was measured by adding HRP-conjugated DDDK tag antibody, (ab1238, abcam; 1:10,000) in ELISA buffer (1% goat serum/ 5% Tween-20/PBS) and perfoming chemiluminescent detection with SuperSignal ELISA Pico chemiluminescent substrate (Pierce). Renilla-HA-NOD2 signal in each well was normalized to matching His-FLAG-CAD measurements, and values were displayed in fold change, relative to no treatment sample.

### Preparation of Cytoplasmic and Nuclear Extracts

RAW-264.7 cells were treated overnight with 200 U/mL IFNβ and 100 μM PALA. Cells were washed and re-suspended in ice-cold PBS. Cells pellets were generated by centrifugation at 1200 RPM for 5 min and re-suspended in buffer A (10 mM HEPES, 1.5 mM MgCl_2_, 1 0mM KCL, 0.5 mM DTT, 0.05% NP40, and 1X protease inhibitor cocktail (Sigma). Nuclei were separated from the cytosol via centrifugation at 1000 x *g* for 10 min at 4°C and washed 3 times with buffer A. Nuclei were then disrupted in buffer B (5 mM HEPES, 1.5 mM MgCl_2_, 0.2 mM EDTA, 0.5 mM DTT, 26% Glycerol) with 4.6 M NaCl for 30 min. Cytoplasmic fractions were prepared by centrifugation at 20,000 x *g* for 20 min at 4°C and supernatants containing cytoplasmic proteins were collected. Protein concentrations were determined using a Bradford assay kit (Thermo Scientific).

### Immunofluorescent confocal microscopy

RAW-264.7 macrophages, as well as WT and NOD2KO BMDMs were seeded on sterilized glass coverslips. The following day, cells were treated for 1 h with 200 U/mL IFNβ and100μM PALA. Coverslips were fixed with ice-cold methanol for 15 min at −20°C and blocked for 1 h in 3% goat serum/0.3% TX-100/PBS. Coverslips were then incubated with IRF1 antibody diluted 1:500 in antibody dilution buffer containing 1% BSA/0.3% TX-100/PBS overnight at 4°C. Coverslips were incubated in anti-rabbit AlexaFluor 568 (Invitrogen) diluted 1:1,000 in antibody dilution buffer for 1 h at room temperature. Coverslips were mounted in Vectashield containing DAPI (Vector Laboratories Inc.) and visualized using a Leica DM2000 microscope and a Leica SP8 confocal microscope. Image quantification of immunofluorescence was performed Image-Pro software (Media Cybernetics, Inc.).

### Statistical analyses

All statistical analyses were performed on GraphPad Prism software (Version 7.02). One-way Analysis of Variance (ANOVA) with post-hoc Tukey’s multiple comparison test or Student’s t-test were used to determine statistical significance. Results are representative of a minimum of three independent experiment performed in triplicate. Data is presented as the mean ± SEM.

## Acknowledgments

We would like to thank Dr. George Stark for the generous gift of cell lines, Dr. Derek Abbott for providing us with bone marrow from RIP2KO mice, Dr. Judy Drazba and Dr. Gauravi Deshpande in the Cleveland Clinic Imaging Core for assistance with confocal microscopy and image analysis, as well as members of the McDonald laboratory for their support. We thank the National Cancer Institute (NCI)/Division of Cancer Treatment and Diagnosis (DCTD)/Developmental Therapeutics Program (DTP) Open Chemical Repository (http://dtp.cancer.org) for providing PALA for our studies. This work was supported by funding from the Office of the Assistant Secretary of Defense for Health Affairs through the Congressionally Directed Medical Research Programs Peer Reviewed Medical Research Program under Award Nos. W81XWH-12-1-0482 (PR110887), W81XWH16-1-0439 (PR150299), and W81XWH-19-1-0488 (PR181846) to C.M., support from the National Institutes of Health (T32GM137868 and 1S10OD019972-01; R21AI153780 to C.M.O.), and philanthropic funds from the McDonald Family Trust. Opinions, interpretations, conclusions, and recommendations are those of the authors and are not necessarily endorsed by the funders.

