## Supplementary material for "N-phosphonacetyl-L-aspartate enhances type I interferon anti-viral responses through activation of non-canonical NOD2 signaling": SI Appendix

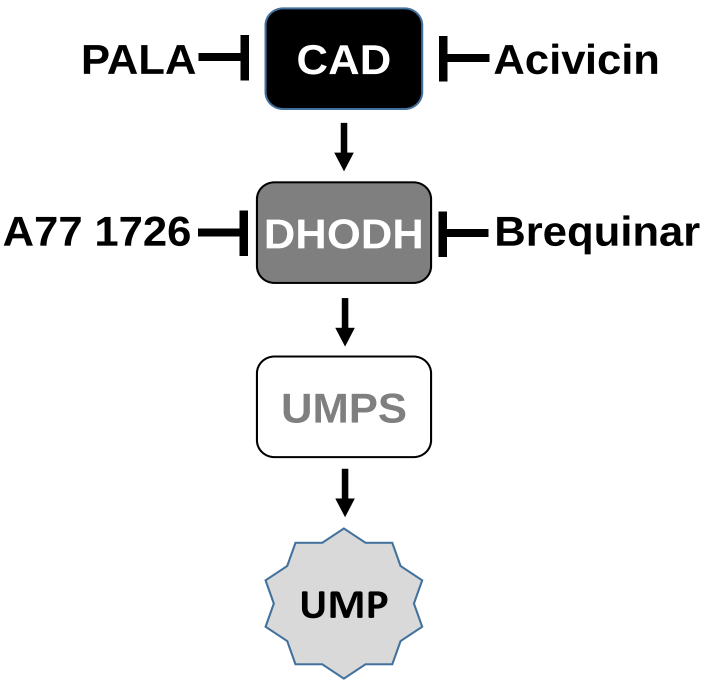


**Figure S1: Illustration of the targets of the *de novo* pyrimidine synthesis inhibitors used in this study.** *N*-phosphonacetyl-L-aspartate (PALA); carbamoyl-phosphate synthetase 2/ aspartate transcarbamylase/ dihydroorotase (CAD); dihydroorotate dehydrogenase (DHODH); uridine monophosphate synthase (UMPS); uridine monophosphate (UMP)

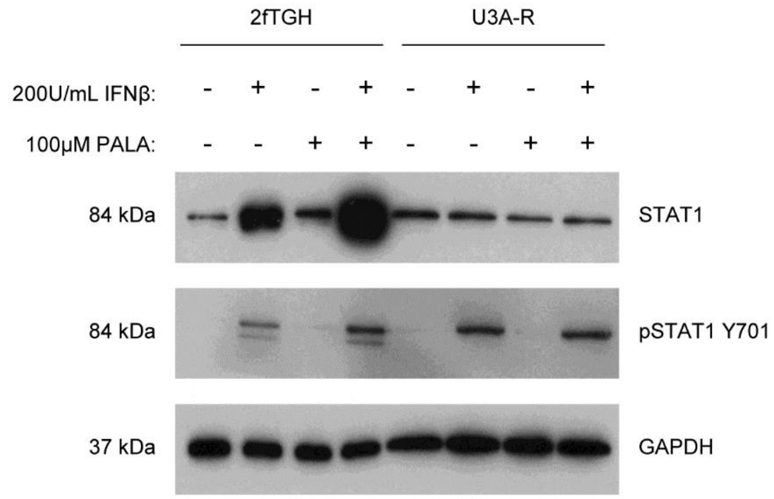


**Figure S2: Enhancement of IFNβ-stimulated STAT1 protein expression by PALA requires the STAT1 promoter.** Immunoblot analysis of STAT1 protein levels in lysates from either cells with endogenous STAT1 expression (2fTGH cells) or a mutant of this cell line that lacks endogenous STAT1 expression but has STAT1 stably expressed from a CMV promoter (U3A-R). Cells were treated for 24h with either IFNβ, PALA, or a combination of IFNβ and PALA. Representative of 3 independent experiments.
